# *De novo* DNA methyltransferase activity in colorectal cancer is directed towards H3K36me3 marked CpG islands

**DOI:** 10.1101/676346

**Authors:** Roza H. Ali Masalmeh, Cristina Rubio-Ramon, Francesca Taglini, Jonathan Higham, Hazel Davidson-Smith, Richard Clark, Jimi Wills, Andrew J. Finch, Lee Murphy, Duncan Sproul

## Abstract

The aberrant gain of DNA methylation at CpG islands (CGIs) is frequently observed in colorectal tumours and may silence the expression of tumour suppressors such as *MLH1*. Current models propose that these CGIs are targeted by *de novo* DNA methyltransferases (DNMTs) in a sequence-specific manner but this has not been tested. Using ectopically integrated CGIs, we find that aberrantly methylated CGIs are subject to low levels of *de novo* DNMT activity in colorectal cancer cells. By delineating DNMT targets, we find that instead *de novo* DNMT activity is targeted primarily to CGIs marked by the histone modification H3K36me3, a mark associated with transcriptional elongation. These H3K36me3 marked CGIs are heavily methylated in colorectal tumours and the normal colon suggesting that *de novo* DNMT activity at CGIs in colorectal cancer is focused on similar targets to normal tissues and not greatly remodelled by tumourigenesis.

## Introduction

DNA methylation is an epigenetic mark associated with gene repression. It is normally pervasive in mammalian genomes but absent from many regulatory elements, particularly CGIs^1^. In tumours, CGIs often become aberrantly methylated. In some cases hypermethylated CGIs correspond to the promoters of tumour suppressor genes such as *MLH1, CDKN2A* (*p16/ARF*) and *BRCA1*^2^. At these genes, hypermethylation associates with repression and thus could drive tumourigenesis. In support of this hypothesis, targeted methylation of the *CDKN2A* promoter in mammary epithelial cells prevents their entry into senescence^3^. However, aberrant CGI hypermethylation also occurs at many other CGIs that are not obviously the promoters of tumour suppressor genes^2^. These aberrantly methylated CGIs are often repressed by Polycomb Repressive Complexes and marked by H3K27me3 in the normal cells that give rise to the cancer^4–6^. However, the mechanisms underpinning the aberrant hypermethylation of CGIs in cancer remain unclear.

DNA methylation is established and maintained in human cells by DNMT1, DNMT3A and DNMT3B^7^. DNMT3A and DNMT3B are *de novo* methyltransferases that establish methylation patterns during early development^8^. DNMT1 is the main enzyme responsible for maintaining DNA methylation^9^. It has a preference for acting at hemimethylated sites *in vitro*^10^ and also interacts with PCNA at replication forks^11^.

DNMT3B is the methyltransferase most often implicated in the aberrant methylation of CGIs in cancer. DNMT3B levels are often increased in cancers relative to normal tissues and higher levels in colorectal tumours correlate with the aberrant methylation of several CGIs^12,13^. Aberrant methylation at CGIs in cancer could be programmed by the sequence-specific recruitment of DNMTs through transcription factors^14^. The transcription factor MAFG is proposed to directly recruit DNMT3B to CGIs methylated in BRAF mutant colorectal tumours^15^. A parallel study suggested that in KRAS mutant colorectal cancer, DNMT1 is recruited by ZNF304 to cause *de novo* methylation of several genes including *CDKN2A*^16^. However, in mouse embryonic stem (ES) cells, DNMT3B is targeted to regions marked by H3K36me3^17^. These observations suggest a model in which DNMT activity in cancer cells is remodelled to target H3K27me3 marked CGIs in a sequence-specific manner resulting in their aberrant methylation. Despite this, previous studies have not measured *de novo* DNMT activity at CGIs in cancer cells and it is unclear whether it is indeed elevated at those that are aberrantly methylated.

Here we use different experimental strategies to assay *de novo* DNMT activity at CGIs in colorectal cancer cells. In contrast to current models, we find that the highest levels of *de novo* DNMT activity are found at CGIs that are marked by H3K36me3 and methylated in the normal colon.

## Results

### CGIs are not *de novo* methylated at ectopic locations in colorectal cancer cells

The ectopic integration of CGIs has been used to demonstrate strong sequencespecific programming of CGI DNA methylation levels by transcription factors in mouse ES cells^18^. Therefore, in order to understand whether *de novo* DNMTs are targeted to CGIs that are aberrantly methylated in colorectal cancer in a sequencespecific manner we asked whether these CGIs become methylated when integrated into ectopic locations in the genome of colorectal cancer cells. If DNMTs were specifically recruited to these CGIs in a sequence-specific manner, the initially unmethylated ectopic integrants would be expected to rapidly acquire high levels of DNA methylation.

We used piggyBac transposons to randomly integrate copies of 10 CGIs into the genome of HCT116 cells (Fig. 1A). We tested 6 aberrantly methylated CGIs that are methylated in HCT116 cells (*CDKN2A, SFRP1, ZFP42, GATA4, CDH7 and CDH13*), a housekeeping CGI promoter that does not become hypermethylated in colorectal cancer, *BUB1*, and a normally methylated CGI, *DAZL*. The CGI promoter of the tumour suppressor gene *MLH1* that is unmethylated in HCT116 cells was also tested. Cells populations carrying CGI integrations were then expanded for 4 weeks to ensure free plasmid was no longer present in the population before we determined DNA methylation levels at the integrated copies and native locus using specific bisulfite PCR primers. Each ectopically integrated copy assayed will derive from a separate integration site in the population of transfected cells thus sampling a diverse array of different genomic locations.

**Figure 1.**
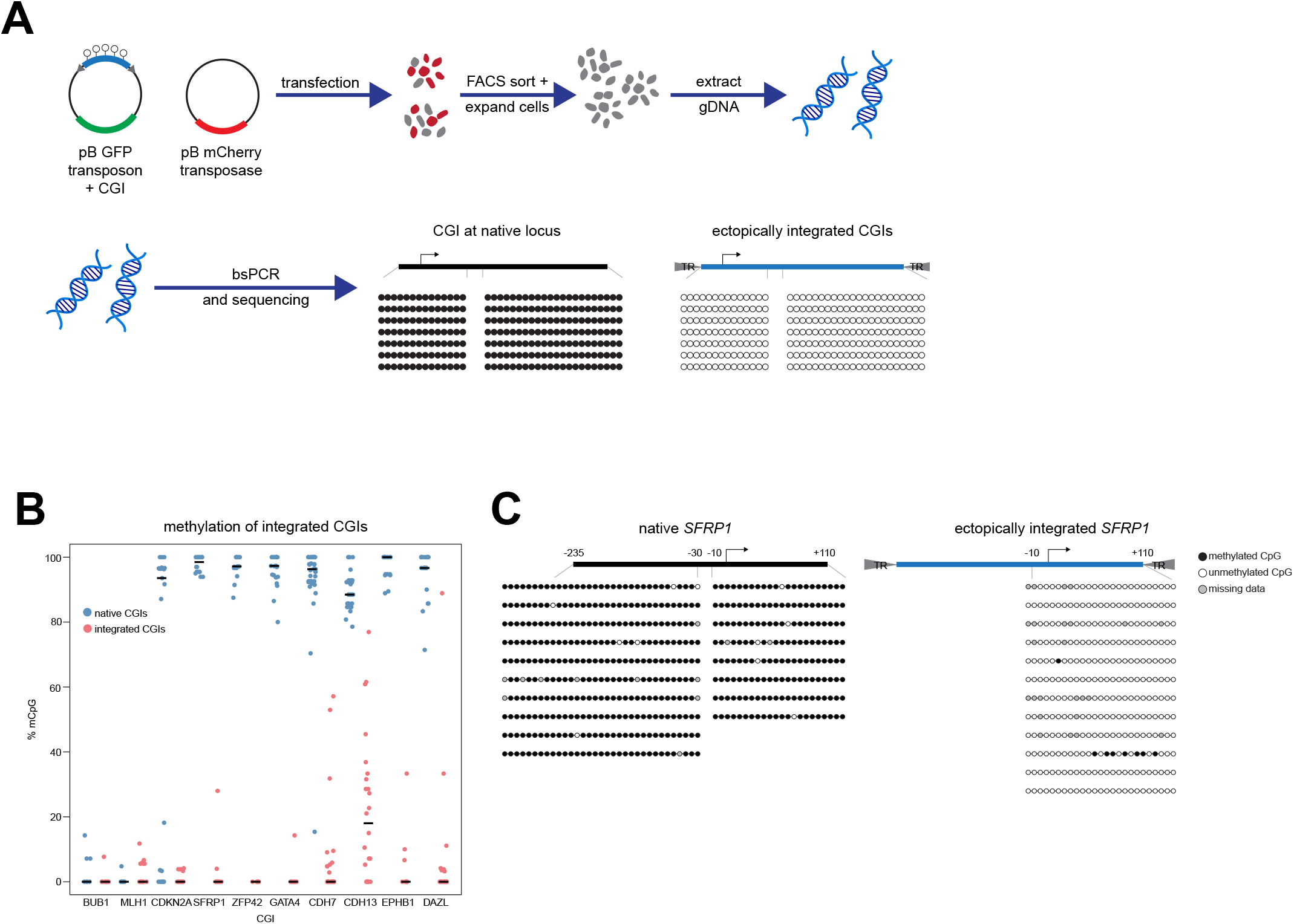
CGIs are not *de novo* methylated at ectopic locations in colorectal cancer. a) Schematic of experimental approach. Cells are transfected with piggyBAC (pB) transposons containing unmethylated CGI DNA sequences along with the pB transposase. Transfected cells are selected by FACS based on fluorescent markers encoded by the plasmids and expanded to dilute out unintegrated copies of the plasmids. Genomic DNA is then extracted and subject to bisulfite PCR (bsPCR) to assay DNA methylation levels at the native and integrated copies of the CGI. b) Ectopically integrated aberrantly methylated CGIs do not become *de novo* methylated in HCT116 cells. Plot showing the mean methylation per clone as assayed by bsPCR for 10 CGIs in their native location and when ectopically integrated in a piggyBac transposon. Each point represents the mean methylation level for a single bsPCR clone. The thick line indicates the median for each CGI. The number of clones analysed per CGI is indicated in Supplementary Table 1. c) *SFRP1* is not hypermethylated when integrated into ectopic locations in HCT116 cells. Illustrative example bsPCR data from panel b showing the *SFRP1* promoter CGI in its native and integrated state. Circles are CpGs with different clones arranged vertically. Each integrated clone derives from a separate genomic integration. Black circles are methylated CpGs and white circles are unmethylated CpGs. Grey circles represent missing data due to sequencing errors.

The *BUB1* and *MLH1* CGIs both remained unmethylated when integrated into ectopic locations using piggyBac (Fig. 1B). The vast majority of integrated copies of the aberrantly methylated CGIs tested also did not become *de novo* methylated (Fig. 1B,C). Rare cases of methylation observed at integrated copies were low level and heterogeneous (Fig. 1C) but confirmed that HCT116 cells are capable of *de novo* methylation as previously reported^19^. The highest levels of methylation observed were at ectopic copies of the *CDH13* CGI but these were still much lower than those seen at the native locus (Fig. 1B). Ectopic copies of the *DAZL* CGI also did not become methylated to a high level (Fig. 1B) suggesting that it is only targeted by *de novo* DNMTs when it gains methylation early in development^20^.

Overall, the results of this experiment suggest that in HCT116 cells surprisingly low *de novo* DNMT activity is targeted in a sequence-specific manner to those CGIs that frequently become hypermethylated in colorectal cancer.

### DNMT3B targets H3K36me3 marked CGIs in colorectal cancer cells

Given that our experiments with ectopically integrated CGIs suggested that aberrantly methylated CGIs were not strongly targeted by *de novo* DNMT activity, we next sought to determine which CGIs were targeted by DNMTs in colorectal cancer. We focused on DNMT3B because of its prior associations with aberrant methylation of CGIs in colorectal cancer^12,13^. We therefore reintroduced DNMT3B2, the major catalytically active DNMT3B isoform expressed in somatic cells (henceforth referred to as DNMT3B)^21^, into hypomethylated HCT116 cells which lack DNMT3B and express low levels of a truncated DNMT1 product (DKO cells)^22,23^. Regions of the genome targeted by DNMT3B will gain methylation in this experiment. DNMT3B expression in DKO cells led to increased total methylation levels, as measured by mass-spectrometry, from 60.6% to 81.1% of the level observed in HCT116 cells (gain of 20.5%, Supplementary Fig. 1A). A gain of 9.04% was also observed when catalytically dead DNMT3B was reintroduced (Supplementary Fig. 1A).

We then used reduced representation bisulfite sequencing (RRBS)^24^ to determine which CGIs gained methylation in DKO cells upon reintroduction of DNMT3B. In this experiment 2,238 CGIs gained significant levels of methylation and are putative DNMT3B targets (Fig. 2A, ≥20% methylation gain and Benjamini-Hochberg corrected Fisher’s exact tests p < 0.05). When cross-referenced to HCT116 histone modification ChIP-seq data from ENCODE^25^, these CGIs were significantly enriched in regions marked by H3K36me3 (Fig. 2B, C). To directly examine the relationship between H3K36me3 and gain of methylation upon DNMT3B expression, we ranked CGIs by their H3K36me3 level in HCT116 cells. Those CGIs with the highest levels of H3K36me3 in HCT116 cells gained the most DNA methylation in DKO when DNMT3B is reintroduced (Fig. 2D).

**Figure 2.**
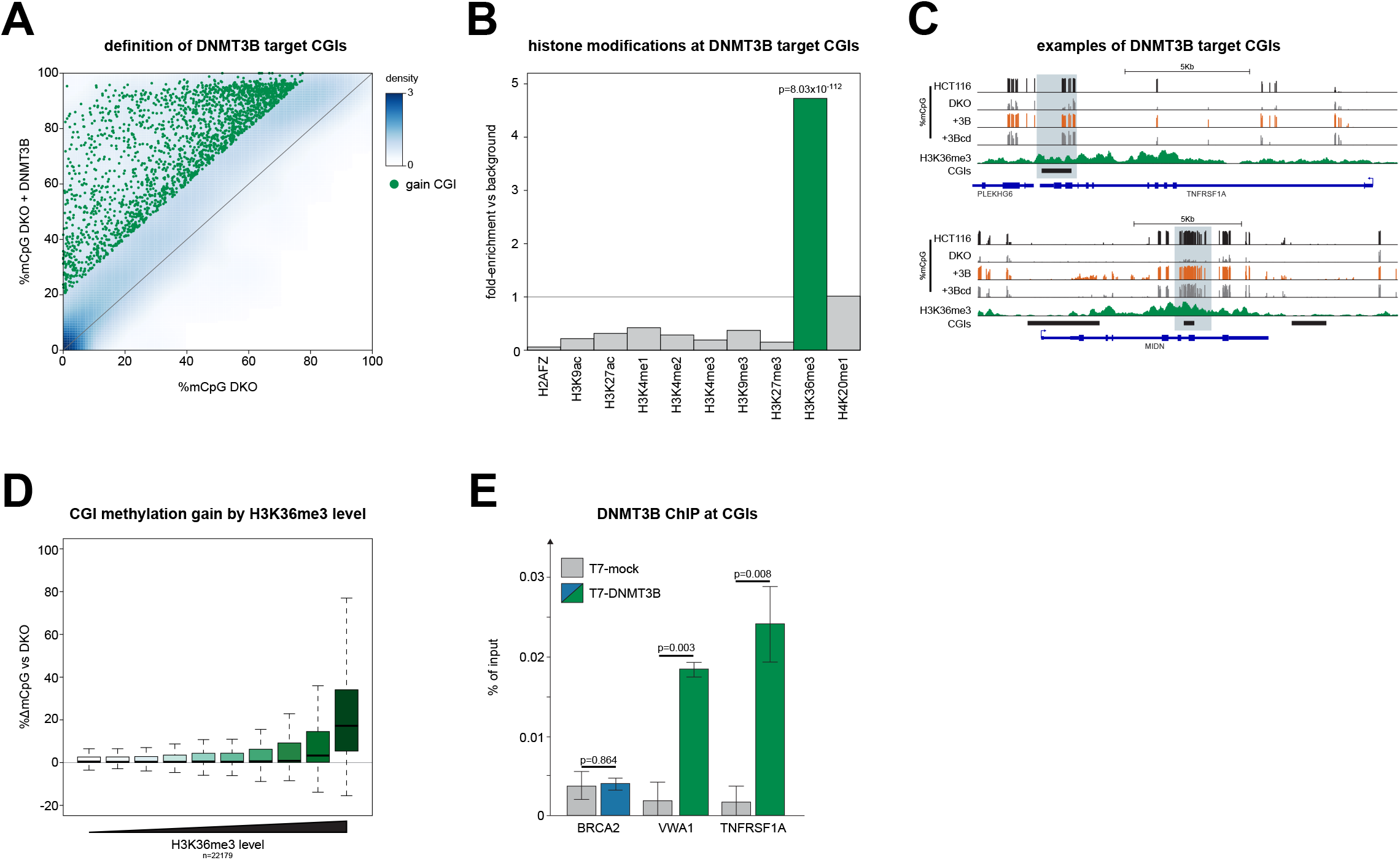
DNMT3B targets H3K36me3 marked CGIs in colorectal cancer cells. a) Many CGIs gain methylation upon expression of DNMT3B in DKO cells. Density scatter plot comparing mean CGI methylation levels in DKO cells and DKO cells expressing DNMT3B (DKO + DNMT3B). Green individual points denote CGIs significantly gaining methylation upon DNMT3B expression (≥20% methylation gain and Benjamini-Hochberg corrected Fisher’s exact tests p < 0.05). b) DNMT3B target CGIs are enriched in H3K36me3 marked regions. Barplot of the fold-enrichment observed for HCT116 histone modification peaks in the set of CGIs that significantly gain methylation when DNMT3B is expressed in DKO cells compared to the background of all CGIs observed in the experiment. P-values are from Fisher’s exact tests (1-sided for enrichment). c) Examples of DNMT3B target CGIs. Genome browser plots showing DNA methylation levels and HCT116 H3K36me3 ChIP signal. CGIs and genes are shown below the plots. CGIs gaining methylation when DNMT3B is expressed in DKO cells are indicated in light blue. +3B = DKO + DNMT3B; +3Bcd = DKO + catalytically dead DNMT3B. Scale for methylation data is 0 to 100%. For H3K36me3 it is 0 to 9.5. d) CGIs with the highest levels of H3K36me3 gain the greatest levels of DNA methylation. Boxplot showing gain of DNA methylation at CGIs when DNMT3B is expressed in DKO cells relative to their level of H3K36me3. CGIs were ranked by H3K36me3 and split into 10 equally sized groups. Lines=median; Box=25th–75th percentile; whiskers=1.5× interquartile range from box. e) DNMT3B localises to CGIs that gain methylation upon DNMT3B expression in DKO cells. qPCR analysis of CGIs following anti-T7 ChIP from T7-DNMT3B cells and HCT116 cells lacking the tag (T7-mock). Shown are the mean of 3 biological replicates with the standard deviation indicated by the error bars. P-values are from T-tests.

We then conducted a second experiment, expressing DNMT3B to a higher level (using the CAG promoter rather than EF-1α, Supplementary Fig. 1B). This resulted in a greater gain of methylation at H3K36me3 marked CGIs and more CGIs were restored to the levels observed in HCT116 cells (Supplementary Fig. 1C, D). Ectopic gains of methylation at loci hypomethylated in HCT116 cells were also observed in this 2^nd^ experiment (Supplementary Fig. 1C, arrow). Catalytically dead DNMT3B also caused a significant gain in DNA methylation at H3K36me3 marked CGIs compared to a GFP expressing control although this was to a lower level than that seen with catalytically active DNMT3B (Supplementary Fig. 1C, D). The level of methylation gain seen with catalytically dead DNMT3B was similar in both the low and high expression experiments (Supplementary Fig. 1D). DNMT3A and DNMT3B are known to interact^26^. Interactions between catalytically dead DNMT3B and DNMT3A could therefore be responsible for these gains as DNMT3A is upregulated in DKO cells compared to HCT116 cells (Supplementary Fig. 1E).

To confirm that DNMT3B normally targets CGIs gaining methylation in our DKO experiments, we used CRISPR to introduce an N-terminal T7 tag on DNMT3B in HCT116 cells (T7-DNMT3B cells) and performed ChIP-qPCR. N-terminal tagged DNMT3B is catalytically active *in vivo*^27^ and we observed no loss of methylation at representative CGIs in T7-DNMT3B cells (Supplementary Fig. 1F). DNMT3B was significantly enriched at gene body CGIs in the *VWA1* and *TNFRSF1A* genes that are methylated in HCT116 and gained methylation when DNMT3B was expressed in DKO cells (Fig. 2E). No enrichment of DNMT3B was seen at the *BRCA2* promoter CGI, a housekeeping gene that does not gain methylation in DKO cells expressing DNMT3B (Fig. 2E). Taken together, our experiments in DKO cells and T7-DNMT3B cells suggest that DNMT3B is primarily targeted to CGIs that are marked by H3K36me3 in colorectal cancer.

### H3K36me3 marked CGIs preferentially recover methylation following pharmacological hypomethylation

To quantify *de novo* DNMT activity at CGIs without ectopically overexpressing or focusing on specific DNMTs, we hypomethylated HCT116 cells using the demethylating drug 5-aza-2’-deoxycytidine (5-aza-dC) before measuring their recovery of DNA methylation. At 3 days following 5-aza-dC treatment, total DNA methylation had decreased to 49.9% of that in untreated cells (Supplementary Fig. 2A). HCT116 cells recovered methylation to levels similar to untreated cells within 22 days after 5-aza-dC treatment (Supplementary Fig. 2A). In order to compare the relative recovery rate of different CGIs, we performed RRBS across this time-course (Fig. 3A). The re-methylation observed in this experiment could be the result of *de novo* DNMT activity or the outgrowth of cells that escaped 5-aza-dC induced hypomethylation. However, we observed significant heterogeneity in the normalised recovery rates for different CGIs (p=1.31×10^−25^ by ANOVA, see materials and methods) suggesting that the recovery of methylation was not solely explained by the outgrowth of cells escaping 5-aza-dC induced hypomethylation.

**Figure 3.**
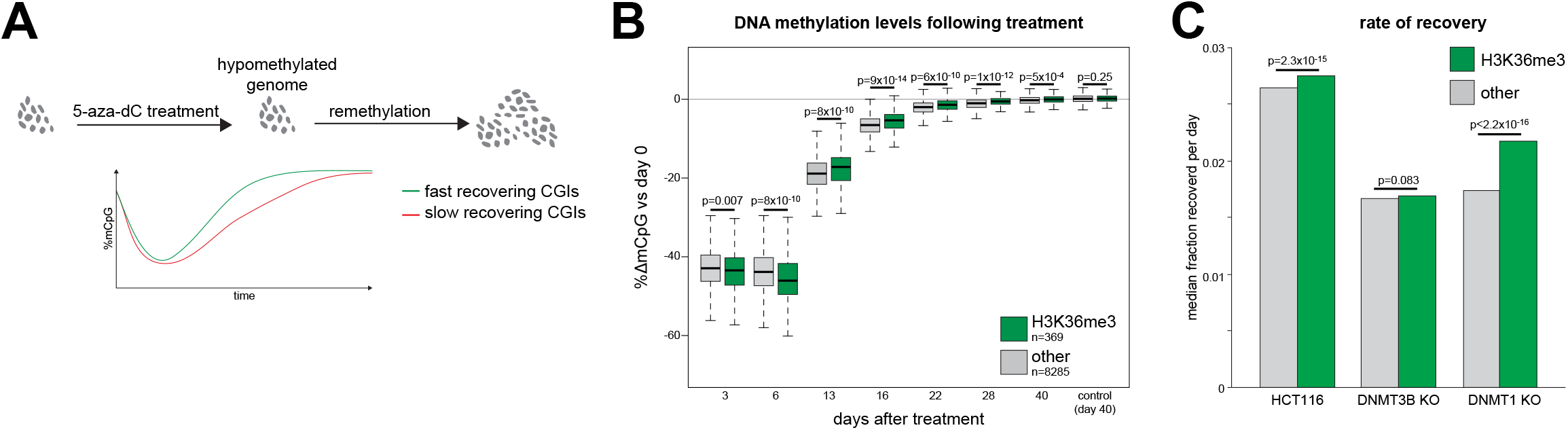
H3K36me3 marked CGIs preferentially recover methylation following pharmacological hypomethylation. a) Schematic of experimental approach. 5-aza-dC is used to hypomethylate cells and the kinetics of methylation recovery are compared for different CGIs. b) H3K36me3 marked CGIs preferentially recover methylation following 5-aza-dC treatment. Boxplots of relative methylation at H3K36me3 marked CGIs and all other CGIs methylated in HCT116 cells. P-values are from Wilcoxon rank sum tests. Lines=median; Box=25th–75th percentile; whiskers=1.5× interquartile range from box. c) The rate of methylation gain is higher at H3K36me3 CGIs in HCT116 cells. Barplot of the median normalised rate of methylation gain for H3K36me3 marked CGIs and other CGIs methylated in HCT116 cells and DNMT KO cells. The rate of methylation recovery was estimated by fitting linear models to the RRBS data for each CGI. P-values are from Wilcoxon rank sum tests.

H3K36me3 marked CGIs lost significantly more methylation than other HCT116 methylated CGIs at 3 and 6 days following 5-aza-dC treatment (Fig. 3B). However, we observed that H3K36me3 marked CGIs recovered DNA methylation significantly more rapidly than other CGIs at later time-points (Fig. 3B). Quantification of the individual normalised rates of methylation gain for each CGI confirmed that remethylation was significantly faster at H3K36me3 marked CGIs than other CGIs (Fig. 3C). We then performed the same experiment in HCT116 cells lacking DNMT3B (DNMT3B KO cells) or DNMT1 (DNMT1 KO cells)^23^. Loss of DNMT3B attenuated the difference in re-methylation rate between H3K36me3 and other CGIs (Fig. 3C, Supplementary Fig. 2B) whereas the difference was exacerbated in DNMT1 KO cells (Fig. 3C, Supplementary Fig. 2C). This is consistent with the difference in remethylation kinetics observed in HCT116 cells being caused by differential *de novo* DNMT activity and suggests that H3K36me3 marked CGIs are preferential targets of *de novo* DNMT activity in colorectal cancer cells.

### H3K36me3 marked CGIs are methylated in colorectal tumours and the normal colon

Given that our experimental data suggested that *de novo* DNMT activity in colorectal cancer cells is primarily targeted to CGIs marked by H3K36me3, we wanted to understand whether H3K36me3 patterns were remodelled in colorectal cancer to cause aberrant hypermethylation at some CGIs.

We defined colorectal tumour H3K36me3 marked CGIs by re-analysing H3K36me3 ChIP-seq data from a colorectal tumour^28^. Using TCGA Infinium methylation array data from 342 colorectal tumours and 42 normal colon samples^29^ we found that the vast majority of H3K36me3 CGI probes were highly methylated in clinical specimens (Fig. 4A-C). In contrast, CGI probes associated with H3K4me3, a mark that repels DNMT3 enzymes^30^, had significantly lower methylation levels in clinical colorectal tumours (Fig. 4B-C).

**Figure 4.**
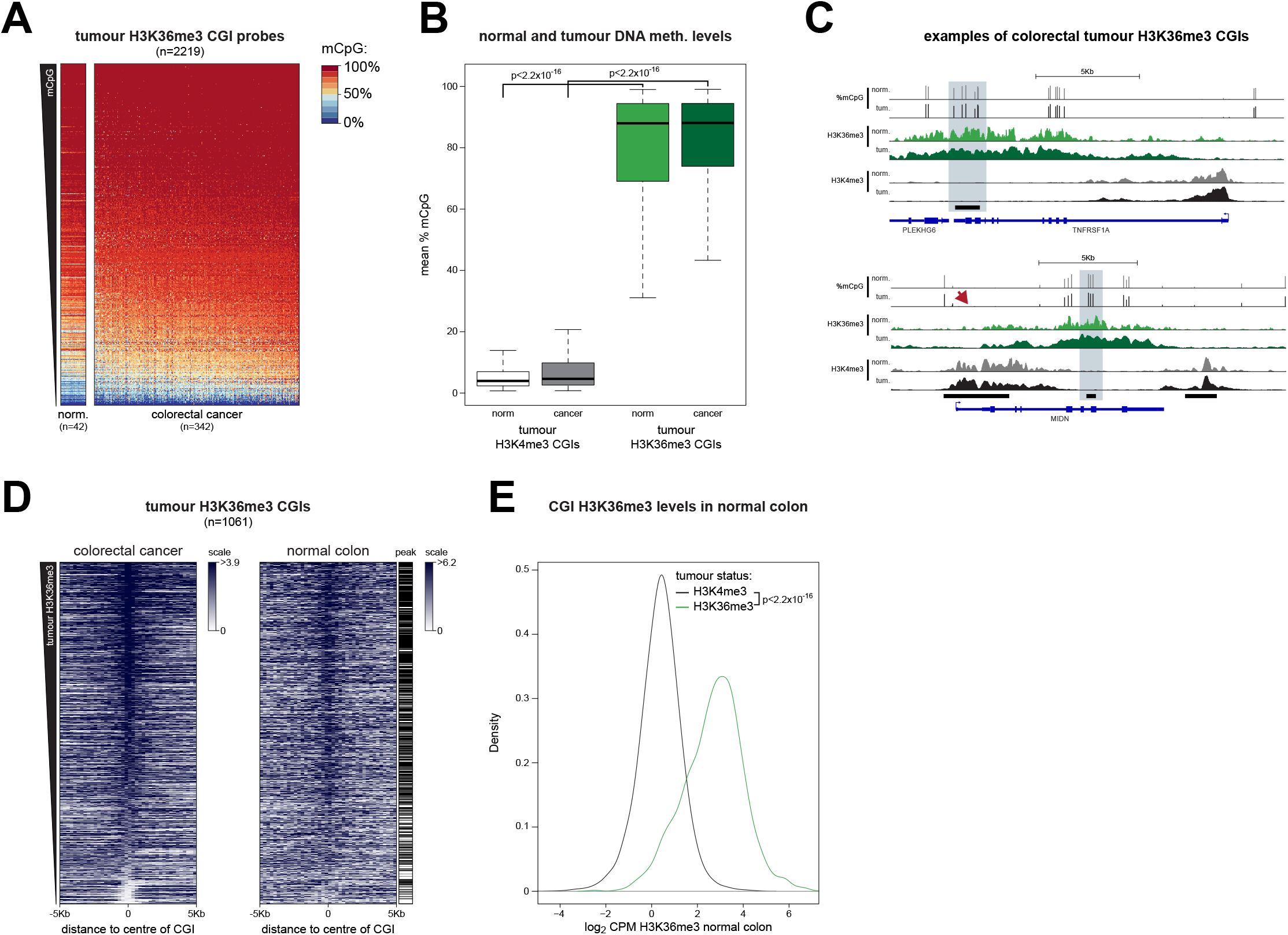
H3K36me3 marked CGIs are methylated in colorectal tumours and the normal colon. a) H3K36me3 marked CGIs are methylated in colorectal tumours. Heatmaps of methylation levels at probes located within H3K36me3 peaks and CGIs in colorectal tumours and adjacent normal colon samples. Both CpGs and samples are ordered by mean DNA methylation levels in colorectal tumours. norm. = normal colon. b) H3K36me3 marked CGIs are more highly methylated in colorectal tumours than H3K4me3 marked CGIs. Boxplot of the mean CpG methylation for probes located within CGIs and H3K36me3 or H3K4me3 peaks for 42 normal colon samples and 342 colorectal tumours. P-values are from Wilcoxon rank sum tests. Lines=median; Box=25th–75th percentile; whiskers=1.5× interquartile range from box. c) Examples of colorectal tumour H3K36me3 marked CGIs. Genome browser plots showing mean DNA methylation levels along with H3K36me3 and H3K4me3 signal for both the normal colon and colorectal tumours. CGIs and genes are shown below the plots. CGIs marked by H3K36me3 in the colorectal tumour are indicated in light blue. Arrow indicates lack of methylation at an H3K4me3 marked CGI. norm. = normal colon, tum. = colorectal tumour. Scale for methylation data is 0 to 100%. d) Colorectal tumour H3K36me3 CGIs are marked by H3K36me3 in the normal colon. Heatmaps of H3K36me3 ChIP-seq at colorectal tumour H3K36me3 CGIs. The left panel shows colorectal tumour H3K36me3 levels and the right panel normal colon H3K36me3 levels at the same CGIs. CGIs are ordered by mean H3K36me3 enrichment in the colorectal tumour. Those overlapping ChIP-seq peaks in the normal colon are indicated next to the heatmap (black=peak). e) Colorectal tumour H3K36me3 CGIs have high levels of H3K36me3 in the normal colon. Density histogram of normalised normal colon H3K36me3 counts at CGIs marked by H3K36me3 and H3K4me3 in the colorectal tumour. P-value from Wilcoxon rank sum test.

Our analysis also demonstrated that H3K36me3 marked CGIs were highly methylated in the normal colon samples (Fig. 4A-C). An analysis of 38 matched tumour-normal pairs from TCGA confirmed that these CGIs had similar methylation levels in the normal colon and colorectal tumours (mean Pearson correlation=0.826). The correlation between matched samples for H3K4me3 peaks was significantly lower (mean 0.694, p=2.90×10^−9^, Wilcoxon rank sum test). Using data from the Roadmap Epigenomics project, we then asked if colorectal tumour H3K36me3 marked CGIs were also associated with H3K36me3 in the normal colon. Our analysis showed strong normal colon H3K36me3 signal at these CGIs and 758/1061 overlapped a normal colon H3K36me3 peak (71.4%, p < 2.2×10^−16^, Fishers exact test, Fig. 4D). In contrast, colorectal tumour H3K4me3 marked CGIs had significantly lower levels of H3K36me3 in the normal colon (p < 2.2×10^−16^, Wilcoxon rank sum test) and were significantly depleted in normal colon H3K36me3 peaks (4.21% overlapped, p < 2.2×10^−16^, Fishers exact test, Fig. 4E)

Taken together, these analyses suggest that the CGIs that are marked by H3K36me3, which are subject to high *de novo* DNMT in colorectal cancer cells, are highly methylated in both clinical colorectal tumours and the corresponding normal tissue and that H3K36me3 patterns are not extensively remodelled at CGIs in colorectal tumours.

## Discussion

Here we present the first comprehensive assessment of *de novo* DNMT activity at CGIs in colorectal cancer. Our results suggest that this is primarily directed towards gene body CGIs marked by H3K36me3, a histone modification associated with ongoing active transcription, and lower levels are targeted to aberrantly methylated CGIs (Fig. 5). We find that these CGIs are not only highly methylated in tumours but also the normal colon suggesting that *de novo* DNMTs target similar CGIs in both normal somatic tissues and cancers.

**Figure 5.**
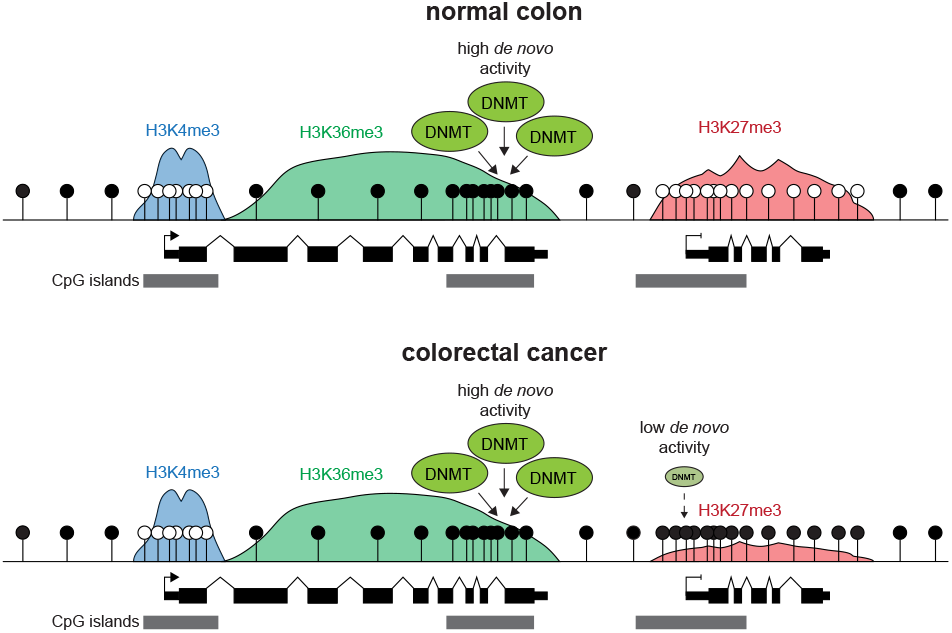
*De novo* DNA methyltransferase activity in colorectal cancer is directed towards H3K36me3 marked CpG islands. Graphical model of the distribution of *de novo* DNMT activity at CGIs in colorectal cancer. Our data support a model whereby the CGIs subject to the highest levels of de novo DNMT activity in both the normal and cancerous colon are those marked by H3K36me3. Although CGIs marked by H3K27me3 in the normal colon become aberrantly methylated in colorectal cancer^4–6^, our results suggest that this is associated with inefficient *de novo* methylation. H3K27me3 is lost from these CGIs when they become aberrantly methylated^60^.

Our observation that the highest levels of *de novo* methylation at CGIs in cancer cells are targeted towards those associated with H3K36me3 parallels observations made in diverse systems. Both DNMT3A and DNMT3B possess a PWWP domain that binds H3K36me3^31^. In mouse embryonic stem cells DNMT3B is primarily localised to H3K36me3^17^ and DNMT3B knockout leads to preferential loss of DNA methylation from H3K36me3 marked regions^32^. H3K36me3 is deposited by SETD2 through its association with RNA polymerase II^33^. Transcription induced deposition of H3K36me3 leads to DNMT3B-dependent methylation of CGIs in mouse ES cells^34^ and is associated with *de novo* methylation of imprinting control regions in mouse oocytes^35^. Previous work in DKO cells suggests DNMT3B is targeted to H3K36me3 in gene bodies but did not examine CGIs^36^.

Here we have focused on assays that detect *de novo* methylation irrespective of the enzyme involved. Overall, they suggest that *de novo* DNMT activity is primarily targeted to H3K36me3 marked CGIs irrespective of the DNMT responsible but support a model in which the bulk of the *de novo* DNMT activity at H3K36me3 marked CGIs is dependent of DNMT3B. We also observe gains of DNA methylation at H3K36me3 marked loci when catalytically inactive DNMT3B is introduced into DKO cells. This correlates with increased DNMT3A levels in DKO cells. DNMT3A and B can interact^26^ and a previous study suggested that catalytically inactive DNMT3B may recruit DNMT3A to H3K36me3 marked gene bodies^36^. Part of the *de novo* DNMT activity we measure at H3K36me3 marked CGIs in colorectal cancer cells could therefore be mediated by DNMT3A rather than DNMT3B but is dependent on DNMT3B for recruitment.

Current models suggest that DNMTs are targeted to aberrantly methylated CGIs in colorectal cancer by transcription factors^15,16^. Aberrantly methylated CGIs are generally repressed in normal cells and associated with H3K27me3 and Polycomb Repressive Complexes rather than H3K36me3^4–6^. Our results in this study suggest that the sequences of aberrantly methylated CGIs inefficiently recruit *de novo* DNMTs and are inconsistent with models of *de novo* methylation based on recruitment by transcription factors. A similar assay has previously been used to demonstrate sequence-specific programming of DNA methylation at CGIs by transcription factors in mouse ES cells^18^. Our observations suggest that aberrant CGI hypermethylation could occur through an inefficient, slow process associated with low *de novo* DNMT activity. In support of this hypothesis, the aberrantly methylated *GSTP1* CGI promoter acquires little methylation when it is ectopically introduced into prostate cancer cells but gains of methylation were stimulated by prior *in vitro* seeding of low-level DNA methylation^37^.

Here we have used an established cancer cell line that does not model early stages of transformation. It is possible that a wave of *de novo* DNMT activity is targeted to aberrantly methylated CGIs specifically at the point of transformation paralleling the wave of genome-wide *de novo* methylation observed during early development^38^. If the signals causing this wave of *de novo* DNMT activity were subsequently lost in advanced tumours, they would not be ascertained in experiments using established cancer cell lines. However, oncogenic mutations are proposed to be the signal responsible for instructing *de novo* methylation^15,16^ and these persist in advanced cancers and cell lines. HCT116 cells also have the ability to *de novo* methylate retroviral DNA sequences^19^ and we observe some *de novo* methylation at ectopic CGIs.

H3K36me3 has not been extensively examined in colorectal cancer, but gains of DNA methylation in cancer have previously been associated with transcription across CGIs. In colorectal and breast tumours the *TFPI2* promoter is aberrantly hypermethylated in association with transcription originating from a nearby LINE-1 promoter^39^. Also Lynch syndrome can be caused by the constitutive hypermethylation of the tumour suppressor *MSH2* associated with read-through transcription from the upstream gene caused by a genetic deletion^40^. Given the association of SETD2 with RNA polymerase II^33^, these cases of transcription-associated hypermethylation would be expected to be associated with H3K36me3 deposition as observed in experiments examining the effect of transcription across CGIs in differentiating mouse ES cells^34^. We find that colorectal tumour H3K36me3 marked CGIs are also associated with H3K36me3 and high levels of DNA methylation in the normal colon. This suggests that H3K36me3 patterns are not extensively remodelled in colorectal tumours compared to normal tissue and that transcription-coupled deposition of H3K36me3 is an infrequent cause of aberrant CGI hypermethylation in colorectal tumours. Instead the recruitment of high *de novo* DNMT activity to H3K36me3-marked gene body CGIs might serve to prevent spurious transcription from these potential promoter sequences interfering with the expression of the associated genes^1,32^.

Taken together our study suggests that the targeting of *de novo* DNMT activity to CGIs in colorectal cancer is similar to that in normal cells and is predominantly centred on CGIs marked by H3K36me3.

## Methods

### Cell culture

HCT116, DNMT1 KO, DNMT3B KO, and DKO cells were gifts from B. Vogelstein^22,23^. Cells were cultured in McCoy’s 5A medium (Gibco) supplemented with 10% fetal calf serum (Life technologies) and penicillin-streptomycin antibiotics at 140 and 400 μg/ml respectively.

### Generation of plasmid constructs

To create piggyBac transposon constructs containing CGIs, we amplified the CGIs from HCT116 cell genomic DNA and cloned them into the pEGFP-N2-pB-min containing piggyBac terminal repeats. To generate the plasmid pEGFP-N2-pB-min we removed the multiple cloning site (MCS) of the pEGFP-N2 plasmid (Clontech) by BamHI-NheI double digestion, Klenow-mediated blunt ending and ligation. We then added a new MCS flanked by the minimal terminal repeats sufficient for PiggyBac transposition^41,42^. The new MCS was generated from two commercially synthesized oligonucleotides (IDT, oligo sequences in Supplementary Table 2) and inserted into the vector by In-Fusion cloning (TakaraBio). CGI sequences were amplified using specific primers with 15bp overhangs (Supplementary Table 2 for primer sequences) to facilitate In-Fusion cloning into pEGFP-N2-pB-min following the manufacturer’s instructions.

To create the piggyBac DNMT3B expression vector, the DNMT3B2 sequence from pcDNA3/Myc-DNMT3B2 plasmid (Addgene plasmid 36942, a gift from A. Riggs)^27^ was subcloned using XbaI and BamHI into the pCG plasmid (a gift from N. Gilbert) downstream of the T7 tag. The catalytically inactive point mutation C265S^43^ was introduced using the QuikChange II site-directed mutagenesis kit (Agilent) to create T7-DNMT3B2 catalytic dead (CD). T7-DNMT3B2 and T7-DNMT3B2-CD were then cloned into pB530A-puroVal2 using BamHI and EcoRI (weak-expression vector with EF-1α promoter), and in PB-CGIP by swapping eGFP using AgeI and NotI (strong expression vector with CAG promoter, a gift from M. McCrew)^44^. pB530A-puroVal2 was previously created by substituting the copepod GFP from the pB530A plasmid (System Biosciences) with a Puromycin resistance gene.

### Ectopic integration of CGIs

HCT116 cells were transfected using FuGENE HD transfection reagent (Promega). To ectopically integrate CGIs, mCherry expressing piggyBac transposase (gift from W. Akhtar)^45^ was co-transfected with piggyBac CGI transposons into HCT116 cells in 1:1 ratio. After 48 hrs, cells expressing both GFP and mCherry were selected by FACS and expanded for 4 weeks to dilute out free plasmid. DNA was purified with phenol– chloroform extraction and ethanol precipitation. The methylation levels of CGIs at native and ectopic locations were then assessed by bisulfite PCR (primers listed in Supplementary Table 2).

### DNMT3B expression in DKO cells

DKO cells were transfected with FuGENE HD. DNMT3B expression constructs were co-transfected with mCherry expressing piggyBac transposase. After 48 hours, cells stably expressing DNMT3B were selected with 1.5 μg/ml puromycin (Thermo Scientific). Cells were expanded in the presence of puromycin for 20 days before genomic DNA was extracted using DNeasy Blood & Tissue Kit (QIAGEN) following the manufacturer’s instructions.

### Bisulfite PCR

500ng genomic DNA was bisulfite converted with the EZ DNA Methylation-Gold kit (Zymo Research) according to the manufacturer’s instructions. PCR was conducted using FastStart PCR Master Mix (Roche), purified using the QIAquick PCR purification kit (QIAGEN) and sub-cloned into pGEMT-easy (Promega). Primer sequences are listed in Supplementary Table 2. Individual positive bacterial colonies were Sanger sequenced using SP6 or T7 sequencing primers (Supplementary Table 2) and analyzed using BIQ Analyzer software (for piggyBAC CGI experiment)^46^ or BISMA (for T7-DNMT3B cell line)^47^. To generate figures, results were then extracted from the BIQ or BISMA output HTML files using custom R scripts. Clones with ≥25% sequencing errors were excluded from analyses.

### 5-aza-2’-deoxycytidine time-course

Cells were plated at ^~^20% confluency. Next day cells were treated with freshly prepared 1μM 5-aza-dC (Sigma Aldrich) for 24 hrs. Cell pellets were collected at different time points following treatment and genomic DNA was extracted using DNeasy Blood & Tissue Kit (QIAGEN) following the manufacturer’s instructions.

### Western blotting

Whole-cell extracts were obtained by sonication in UTB buffer (8 M urea, 50 mM Tris, pH 7.5, 150 mM β-mercaptoethanol) and quantified using A_280_ from a NanoDrop (Thermo Scientific). 40-50μg of protein was analysed by SDS–polyacrylamide gel electrophoresis (SDS–PAGE) using 3-8% NuPAGE Tris-Acetate protein gels (Life Technologies) and transferred onto nitrocellulose membrane using Xcell Sure Lock Mini Cell electrophoresis tanks (Novex) in 2.5mM tris-base, 19.2mM glycine and 20% methanol. Immunoblotting was performed following blocking in 10% Western blocking reagent (Roche) using antibodies against DNMT3B (Cell Signalling Technology, D7O7O, 1:1000), GAPDH (Cell Signalling Technology, 14C10, 1:1000) and DNMT3A (Cell Signalling Technology, 2160, 1:500). Images were acquired with ImageQuant LAS 4000 following incubation with HRP conjugated goat anti-rabbit IgG (Invitrogen, A16110, 1:1000).

### Generation of CRISPR/Cas9-edited HCT116 cell line

A guide RNA was designed using the CRISPR design web tool (http://crispr.mit.edu/) with the corresponding oligonucleotide cloned into pSpCas9(BB)-2A-GFP (pX458, Addgene Plasmid 48138, a gift of F. Zhang). The donor template for homology-directed repair was created by amplifying 3 single stranded DNA oligonucleotides (IDT Ultramers). The external sequences are 125bp homology arms flanking each side of DNMT3B starting codon; the internal oligonucleotide is a triple T7 tag sequence. The amplified donor template was then sub-cloned into pGEMT-easy (Promega). The vectors containing the gRNA sequence and the donor template were co-transfected using FuGENE HD transfection reagent (Promega). GFP +ve cells were selected by FACS 48 h after transfection and plated at clonal density. Individual colonies were grown up, screened by PCR and the positive clone validated by Sanger sequencing. The positive clone carries one copy of DNMT3B with and N-terminal triple-T7-tag. The second DNMT3B allele has been disrupted by the integration of a portion of the pSpCas9(BB)-2A-GFP plasmid downstream the 4th codon of DNMT3B. This is predicted to cause a frameshift with a severely truncated product on the second allele or be subject to nonsense mediated decay. Primers and oligonucleotides used are listed in Supplementary Table 2.

### Chromatin immunoprecipitation and qPCR

1×10^7^ cells were harvested, washed and crosslinked with 1% methanol-free formaldehyde in PBS for 8 min at room temperature. Crosslinked cells were lysed for 10 min on ice in 50 μl of lysis buffer (50mM Tris-HCl pH8, 150mM NaCl, 1mM EDTA, 1% SDS) freshly supplemented with proteinase inhibitor (Sigma-Aldrich). IP dilution buffer (20 mM Tris-HCl pH8, 150 mM NaCl, 1 mM EDTA, 0.1% Triton X-100) freshly supplemented with proteinase inhibitor, DTT and PMSF was added to the samples to reach a final volume of 500μl. As prolonged sonication caused T7-DNMT3B degradation, chromatin was fragmented using Benzonase as recommended by Pchelintsev *et al*.^48^. Briefly, samples were sonicated on ice with Soniprep 150 twice for 30 sec to break up nuclei. Then 200U of Benzonase Nuclease (Sigma) and MgCl2 (final concentration 2.5 mM) were added and samples were incubated on ice for 15 min. The reaction was blocked by adding 10μl of 0.5M EDTA pH 8. Following centrifugation for 30 min at 14,000 rpm at 4 °C, supernatants were collected and supplemented with Triton X-100 (final concentration 1%) and 5% input aliquots were retained for later use. Protein A dynabeads (Invitrogen) previously coupled with T7-Tag antibody (D9E1X, Cell Signalling) in blocking solution (1xPBS, 0.5% BSA) were added and the samples incubated overnight under rotation at 4 °C. Beads were then washed for 10 min at 4 °C with the following buffers: IP dilution buffer 1% Triton X-100 (20 mM Tris-HCl pH 8, 150 mM NaCl, 2 mM EDTA, 1% Triton X-100), buffer A (50 mM Hepes pH 7.9, 500 mM NaCl, 1 mM EDTA, 1% Triton X-100, 0.1% Na-deoxycholate, 0.1% SDS), buffer B (20 mM Tris pH 8, 1mM EDTA, 250 mM LiCl, 0.5% NP-40, 0.5% Na-deoxycholate), TE buffer (1 mM EDTA pH 8, 10 mM Tris pH 8). Chromatin was eluted by incubating the beads in extraction buffer (0.1M NaHCO_3_, 1% SDS) for 15 min at 37 °C. To reverse the cross-linking Tris-HCl pH 6.8 and NaCl were added to a final concentration of 130 mM and 300mM respectively, and IP samples were incubated at 65 °C for 2h. Samples were then incubated at 55 °C for 1h after addition of 40 μg of Proteinase K (Roche) and, finally, 2μl of RNase Cocktail Enzyme Mix (Ambion) were added, following by 1h incubation at 37°C. Input material was similarly de-crosslinked. Samples were purified with the MinElute PCR purification kit (QIAGEN). qPCR was performed using Lightcycler 480 Sybr Green Master (Roche) on a Light Cycler 480 II instrument (Roche). Primers used are listed in Supplementary Table 2. Enrichment was calculated compared to input DNA. Delta Ct was calculated using input Ct values adjusted to 100% and assuming primer efficiency of 2.

### Global measurement of DNA methylation by mass-spectrometry

1μg genomic DNA was denatured at 95°C for 10 min in 17.5μL water. DNA was then digested to nucleotides overnight at 37°C with T7 DNA polymerase (Thermo Scientific). The reaction was inactivated by incubating at 75°C for 10 min. Samples were then centrifuged for 45min at > 12,000*g* and the supernatant transferred into new tubes for analysis. Enzyme was removed by solvent precipitation. The samples were adjusted back to initial aqueous condition and volume and LC-MS was performed on a Dionex Ultimate 3000 BioRS / Thermo Q Exactive system, using a Hypercarb 3 μm x 1 mm x 30 mm Column (Thermo 35003-031030) and gradient from 20 mM ammonium carbonate to 2 mM ammonium carbonate 90% acetonitrile in 5 minutes. Data were acquired in negative mode, scanning at 70k resolution from 300 to 350 m/z. Extracted ion chromatograms were analysed using Xcalibur (Thermo Scientific) to extract peak intensities at the m/z values expected for each nucleotide (based on annotation from METLIN)^49^ following manual inspection to ensure that they were resolved as clear single peaks. The % of 5-methylcytosine present in the sample was calculated as the ratio of the area under the 5-methylcytosine peak to the area under the guanine peak.

### Statistical analysis

Statistical testing was performed using R v3.4.2. All tests were two-sided, unless otherwise stated. Further details of specific analyses provided in the relevant methods sections below.

### CGI definition

CGI annotation was taken from Illingworth *et al*.^50^. Overlapping CGI intervals were merged using BEDtools (*version 2.27.1*)^51^ before they were converted to hg38 positions using the UCSC browser liftover tool. Non-autosomal CGIs were excluded from the analysis as were CGIs located in ENCODE blacklist regions^25^.

### RRBS data generation and processing

Genomic DNA was extracted from cells using DNeasy Blood & Tissue Kit according to the manufacturer’s protocol with some modifications. RNase A/T1 Cocktail (Ambion AM2286) was added to proteinase K and samples were incubated at 37°C for 1hr to remove RNA. At the final step, DNA was eluted with ddH2O instead of AE buffer. DNA was quantified by Nanodrop and Qubit.

200ng of purified DNA samples were processed using the Ovation RRBS Methyl-Seq system kit (NuGen Technologies) according to instructions with modifications. Briefly, 0.5ng unmethylated phage λ DNA (NEB) was spiked into each sample to allow assessment of bisulfite conversion efficiency. The methylation-insensitive restriction enzyme MspI was then used to digest the genomic DNA, and digested fragments were ligated to adapters. Adapter-ligated fragments were then repaired before bisulfite conversion with the Qiagen Epitect Fast Bisulfite Conversion kit. Bisulfite-treated adapter-ligated fragments were amplified by 9 cycles of PCR and purified using Agencourt RNAClean XP beads. Libraries were quantified using the Qubit dsDNA HS assay and assessed for size and quality using the Agilent Bioanalyzer DNA HS kit.

Sequencing was performed using the NextSeq 500/550 high-output version 2 kit (75bp paired end reads) on the Illumina NextSeq 550 platform. As instructed for the NuGen RRBS kit, 12bp index reads were generated to sequence the Unique Molecular Identifiers (UMI) in addition to the index present in the adaptors. Libraries were combined into equimolar pools and run on a single flow cell. 10% PhiX control library (Illumina v3 control library) was spiked in to facilitate sequencing by generating additional sequence diversity. Library preparation and sequencing was performed by the Edinburgh Clinical Research Facility.

Raw Illumina sequencing output from the NextSeq (*bcl* files) were converted to paired FASTQ files without demultiplexing using *bcl2fastq* and default settings (*v2.17.1.14*). These FASTQ files were then demultiplexed using custom python scripts considering indexes with perfect matches to the sample indexes. The different lanes for each sample were then combined.

Sequencing quality was assessed with FASTQC (*v0.11.4*). Low quality reads and remaining adaptors were removed using TrimGalore (*v0.4.1*, Settings: *--adapter AGATCGGAAGAGC --adapter2 AAATCAAAAAAAC*). NuGen adaptors contain extra diversity bases to facilitate sequencing. These were removed using the *trimRRBSdiversityAdaptCustomers.py* Python script provided by NuGen (*v1.11*). The paired end reads were then aligned to the hg38 genome using Bismark (*v0.16.3* with Bowtie2 *v2.2.6* and settings: *-N 0 -L 20*)^52,53^ before PCR duplicates were identified and removed using the 6bp UMIs present in the index reads and the *nudup.py* Python script supplied by NuGen (*v2.3*). Aligned BAM files were processed to report coverage and number of methylated reads for each CpG observed. Forward and reverse strands were combined using Bismark’s methylation extractor and bismark2bedgraph modules with custom Python and AWK scripts. Custom scripts are available from the authors on request.

Processed RRBS files were assessed for conversion efficiency based on the proportion of methylated reads mapping to the λ genome spike-in (>99.5% in all cases). For summary of RRBS alignment statistics see Supplementary Table 3. BigWigs were generated from RRBS data using CpGs with coverage ≥5.

### Analysis of RRBS data

CGI methylation levels were calculated as the weighted mean methylation using the observed coverage of CpGs within the CGI (unconverted coverage/total coverage). Only CpGs with a total coverage ≥ 10 in all samples were considered in each analysis. Methylated CGIs were defined as those where the weighted-mean % methylation was ≥ 70%. CGIs significantly gaining methylation upon expression of DNMT3B in DKO cells were defined as those with a Benjamini Hochberg adjusted Fisher’s exact test p-value < 0.05 and where ≥ 20% methylation was gained. To analyse histone marks at these CGIs, the number overlapping HCT116 ChIP-seq peaks was compared to the number overlapping all CGIs in the analysis using Fisher’s exact tests.

### Analysis of re-methylation kinetics following 5-aza-2’-deoxycytidine treatment

If re-methylation following 5-aza-dC treatment was entirely driven by outgrowth of unaffected cells, we expect the methylation trajectory of each CGI to be proportional to its initial methylation level (Supplementary Fig. 3). In order to test this possibility, we normalised HCT116 methylation data for all CGIs with day 0 mean CGI methylation > 50% over the methylation value at day 0. Two models were fitted to this normalised data using time-points between day 3 and day 22 inclusive. The null hypothesis that re-methylation is due to outgrowth was represented by the mixed linear model shown in equation 1. The hypothesis that each CGI is remethylated at a different rate was represented by a mixed linear model accounting for a different trajectory for each CGI shown in equation 2.

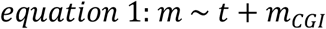

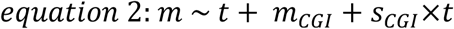

Where *m* is mean normalised methylation, *t* is time, *m_CGI_* is the random intercept for each CGI, and *s_CGI_* is the random slope for each CGI. Comparing the models with ANOVA reveals that the probability that the null hypothesis is a better fit to the data is p=1.31×10^−25^, strongly suggesting that the CGIs do not recover methylation at the same rate. The R function *lmer* from the *lme4* package (*v1.1*) was used for this analysis.

In order to investigate the trajectories of each CGI independently from one another, a linear model was fitted to the normalised methylation of each individual CGI at time points between day 3 and day 22 inclusive. The slopes of these individual linear models are equal to the fraction of methylation recovered per day. This analysis was repeated for the DNMT3B KO and DNMT1 KO cell lines.

### Processing of ChIP-seq data

Processed ChIP-seq data from the normal colon and HCT116 cells were downloaded from ENCODE^25^ as aligned BAM files and replicated peak BED files. The corresponding annotated input control was also downloaded for each sample. bigWig files for visualisation were downloaded from ENCODE. Data for colorectal tumour H3K36me3 and H3K4me3 ChIP-seq^28^ was downloaded as FASTQ files from ENA. Read quality was analysed using FASTQC (*v0.11.4, https://www.bioinformatics.babraham.ac.uk/projects/fastqc*), with low quality reads and adaptors removed using trim-galore with default settings (*version 0.4.1*). Reads were aligned to hg38 using bowtie 2 (*v2.3.1*, with settings: *-N 1 -L 20 --no-unal*)^52^. Multi-mapping reads excluded using SAMtools (*v1.6*, with settings: -bq 10)^54^ and PCR duplicates excluded using SAMBAMBA (*v0.5.9*)^55^.

Tracks for data visualisation were generated using Homer (*v4.8*)^56^. Aligned BAMs were converted to tag directories setting the fragment length to 180bp and converted to bigWig files using Homer’s makeUCSCfile function after filtering with the *removeOutOfBounds.pl* function. ChIP-seq peaks were called from tag directories using Homer’s findPeaks function (settings: *-style histone*) and an input chromatin control sample.

### Analysis of ChIP-seq pileups

Non-overlapping windows of 250bp were defined centred on the midpoint of each CGI, with ChIP-seq read counts/window calculated using BEDtools’ coverage function. Read counts were scaled to counts per 10 million based on total number of mapped reads/sample and divided by the input read count to provide a normalised read count. To prevent windows with zero reads in the input sample generating a normalised count of infinity, an offset of 0.5 was added to all windows prior to scaling and input normalisation. Regions where coverage was 0 in all samples were removed from the analysis. Colour scales for ChIP-seq heatmaps range from the minimum to the 90% quantile of the normalised read count. ChIP-seq peaks were overlapped with CGIs using BEDtools intersect and tested for statistical enrichment with Fisher’s exact tests versus all CGIs included in each analysis. To statistically test differences in histone modification levels, normalised read depths across CGIs were compared using a Wilcoxon rank sum test.

### Infinium array data processing

All available colorectal tumour and normal colon data were downloaded from TCGA Genomic Data Commons as raw IDAT files on 13/9/18^29^. These were processed using the ssNoob approach from the Bioconductor package *minfi* (v*1.22.1*) to derive beta values and detection p-values (beta threshold = 0.001)^57,58^. Individual beta values were excluded where detection p-value was > 0.01. Non-CG probes were also excluded from the analysis. Samples that did not represent primary tumour or adjacent normal colon tissue were excluded using TCGA sample type codes as were samples where ≥ 1% of probes failed the detection p-value threshold. Where replicates existed for these tumour and normal samples, the mean was calculated for each probe. This left 342 tumour samples and 42 adjacent normal colon samples and included 38 matched tumour-normal pairs. Infinium probe locations in the hg38 genome build were taken from Zhou *et al*.^59^. Probes categorised as overlapping common SNPs or having ambiguous genome mapping in that paper were excluded from the analysis (Zhou *et al*. general masking annotation).

### Data availability

All sequencing data that were generated during this study will be deposited in GEO.

## Acknowledgements

We thank I. Kafetzopoulos, W. Bickmore, R. Meehan, P. Heyn, R. Illingworth, C. Tufarelli and G. Ficz for helpful discussions about the study and the Edinburgh Clinical Research Facility Genetics Core and the IGMM mass-spectrometry facility for technical support. This work has made use of the resources provided by the University of Edinburgh digital research services and the MRC IGMM compute cluster. D.S. is a Cancer Research UK Career Development fellow (reference C47648/A20837), and work in his laboratory is also supported by a MRC university grant to the MRC Human Genetics Unit. R.H.A.M. was funded by a HESPAL PhD scholarship from the British Council.

## Contributions

R.H.A.M., C.R.R., F.T. and H.D.S. performed the experiments included in the manuscript. J.W. conducted mass spectrometry supported by A.J.F. and R.C. conducted high-throughput sequencing supported by L.M.. R.H.A.M., J.H. and D.S. conducted the computational analyses of DNA methylation and ChIP-seq. D.S. planned and supervised the study and wrote the manuscript with input from R.H.A.M. and review by all authors.

