## Supplementary tables and figures for "*De novo* DNA methyltransferase activity in colorectal cancer is directed towards H3K36me3 marked CpG islands"

##### Supplementary Table 1

*Clone statistics for bisulphite PCR analysis of CGIs at native and ectopic loci.*

| CGI | Total number of clones | Number of clones by PCR amplicon |
|---|---|---|
| *Native locations* | | |
| *BUB1* | 7 | L: 3, R: 4 |
| *MLH1* | 7 | L: 7 |
| *CDKN2A* | 23 | L: 21, R:2 |
| *SFRP1* | 18 | L: 10, R: 8 |
| *ZFP42* | 16 | L: 10, R: 6 |
| *GATA4* | 25 | L: 16, R: 9 |
| *CDH7* | 41 | L: 19, R1: 17, R2: 5 |
| *CDH13* | 30 | L: 11, R1: 3, R2: 16 |
| *EPHB1* | 20 | R: 20 |
| *DAZL* | 25 | L: 9, R: 16 |
| *Ectopic locations* | | |
| *BUB1* | 10 | L1: 10 |
| *MLH1* | 15 | L: 6, R: 9 |
| *CDKN2A* | 13 | L: 6, R: 7 |
| *SFRP1* | 12 | R1: 6, R2: 6 |
| *ZFP42* | 7 | L: 2, R:5 |
| *GATA4* | 8 | L1: 3, L2: 3, R: 2 |
| *CDH7* | 34 | L: 4, R1: 14, R2: 16 |
| *CDH13* | 24 | L: 7, R1: 10, R2: 7 |
| *EPHB1* | 11 | L: 1, R1: 4, R2: 6 |
| *DAZL* | 22 | L: 8, R: 14 |

##### Supplementary Table 2

*Oligonucleotides used in this study – supplied as an excel spreadsheet.*

##### Supplementary Table 3

*Summary of sequencing statistics for RRBS. Aligned reads counts are following PCR duplicate removal.*

| Sample | Total Reads ($x10^6$) | Aligned Reads ($x10^6$) | Mean CG Coverage | Bisulfite Conversion Rate |
|---|---|---|---|---|
| *DNMT3B expression in DKO cells experiment 1* | | | | |
| HCT116 | 57.98 | 31.22 | 26.83 | 99.67% |
| DKO | 56.16 | 30.61 | 26.53 | 99.69% |
| DKO + DNMT3B | 56.12 | 32.21 | 29.10 | 99.66% |
| DKO + DNMT3Bcd | 52.63 | 28.31 | 25.19 | 99.62% |
| *DNMT3B expression in DKO cells experiment 2* | | | | |
| HCT116 | 38.46 | 22.68 | 20.30 | 99.67% |
| DKO | 42.46 | 25.46 | 23.03 | 99.68% |
| DKO + DNMT3B | 46.99 | 28.40 | 25.83 | 99.68% |

| | | | | |
|---|---|---|---|---|
| DKO + DNMT3Bcd | 46.42 | 27.87 | 25.42 | 99.68% |
| DKO + GFP | 80.56 | 48.64 | 42.78 | 99.68% |
| *5-aza-dC treatment of HCT116* | | | | |
| HCT116 day 0 | As HCT116 in DKO experiment 1 | | | |
| HCT116 day 3 | 60.98 | 30.80 | 25.59 | 99.65% |
| HCT116 day 6 | 50.83 | 26.75 | 23.17 | 99.68% |
| HCT116 day 13 | 49.68 | 28.41 | 25.50 | 99.65% |
| HCT116 day 16 | 61.50 | 32.86 | 27.84 | 99.66% |
| HCT116 day 22 | 52.88 | 28.29 | 24.21 | 99.67% |
| HCT116 day 28 | 60.14 | 33.91 | 30.37 | 99.67% |
| HCT116 day 40 | 53.64 | 28.12 | 23.94 | 99.67% |
| HCT116 day 40 control | 47.91 | 25.66 | 21.76 | 99.67% |
| *5-aza-dC treatment of DNMT3B KO* | | | | |
| DNMT3B KO day 0 | 48.41 | 27.88 | 25.04 | 99.66% |
| DNMT3B KO day 3 | 47.92 | 26.99 | 23.64 | 99.67% |
| DNMT3B KO day 6 | 45.87 | 23.22 | 20.43 | 99.68% |
| DNMT3B KO day 13 | 59.87 | 31.94 | 26.78 | 99.67% |
| DNMT3B KO day 16 | 49.27 | 27.42 | 23.96 | 99.69% |
| DNMT3B KO day 22 | 53.32 | 29.40 | 25.45 | 99.69% |
| DNMT3B KO day 28 | 51.53 | 28.62 | 24.49 | 99.67% |
| DNMT3B KO day 40 | 47.75 | 26.36 | 23.25 | 99.70% |
| DNMT3B KO day 40 control | 54.05 | 30.80 | 27.42 | 99.69% |
| *5-aza-dC treatment of DNMT1 KO* | | | | |
| DNMT1 KO day 0 | 61.57 | 33.50 | 28.23 | 99.66% |
| DNMT1 KO day 3 | 51.79 | 26.85 | 23.45 | 99.68% |
| DNMT1 KO day 6 | 48.56 | 25.17 | 22.35 | 99.67% |
| DNMT1 KO day 13 | 56.15 | 29.72 | 25.92 | 99.68% |
| DNMT1 KO day 16 | 49.88 | 27.04 | 23.28 | 99.66% |
| DNMT1 KO day 22 | 59.70 | 32.34 | 27.56 | 99.67% |
| DNMT1 KO day 40 | 52.22 | 27.10 | 23.67 | 99.68% |
| DNMT1 KO day 40 control | 55.88 | 31.63 | 28.30 | 99.67% |

**Supplementary Figure 1**

**DNMT3B targets H3K36me3 marked CGIs in colorectal cancer cells**

a) Expression of DNMT3B in DKO cells results in a global gain of DNA methylation. Barplot of total methylated cytosine levels estimated by mass-spectrometry. Shown are mean methylation levels relative to HCT116 cells from 3 technical replicates with error bars representing the standard deviation. P-values are from t-tests. +3B = DKO + DNMT3B; +3Bcd = DKO + catalytically dead DNMT3B.

b) Western blots comparing DNMT3B expression levels in two experiments. The upper panels show data from the first experiment (data in *Figure 1*) where the EF-1α promoter drives DNMT3B expression. The lower panels show data from the second experiment where the CAG promoter drives higher levels of DNMT3B. Arrows indicate DNMT3B. +3B = DKO + DNMT3B; +3Bcd = DKO + catalytically dead DNMT3B; +GFP = DKO + GFP.

c) Examples of DNMT3B target CGIs from the second experiment. Genome browser plots showing DNA methylation levels and HCT116 H3K36me3 ChIP signal. CGIs and genes are shown below the plots. CGIs gaining methylation when DNMT3B is expressed in DKO cells are indicated in light blue. Arrow in lower panel indicates a region ectopically gaining methylation with the higher level of DNMT3B expression. +3B = DKO + DNMT3B; +3Bcd = DKO + catalytically dead DNMT3B; +GFP = DKO + GFP. Scale for methylation data is 0 to 100%. For H3K36me3 it is 0 to 9.5.

d) H3K36me3 marked CGIs gain methylation when DNMT3B is expressed in DKO cells. Left, heatmaps of relative methylation levels at methylated HCT116 H3K36me3 marked CGIs. Values denote the change in methylation relative to DKO cells. CGIs are ranked by their mean gain of methylation in experiment 1. Left, boxplots of relative methylation at H3K36me3 marked CGIs and all other CGIs methylated in HCT116 cells. P-values are from Wilcoxon rank sum tests. Lines=median; Box=25th–75th percentile; whiskers=1.5× interquartile range from box.

e) DNMT3A is upregulated in DNMT knockout cells. Western blot for DNMT3A in HCT116 cells, DKO cells and HCT116 cells lacking either DNMT1 (DNMT1 KO) or DNMT3B (DNMT3B KO).

f) T7 tagging of endogenous DNMT3B does not result in losses of DNA methylation at CGIs. Left; barplot showing mean methylation levels estimated by bisulfite PCR at CGIs in HCT116 and T7-DNMT3B cells. Right, representative example bsPCR data from the *TNFRSF1A* gene CGI. Circles are CpGs with different clones arranged vertically. Black circles are methylated CpGs and white circles are unmethylated CpGs.

**Supplementary Figure 2**

**H3K36me3 marked CGIs preferentially recover methylation following pharmacological hypomethylation**

a) HCT116 DNA methylation levels recover 22 days after 5-aza-dC treatment. Barplot of total methylated cytosine levels estimated by mass-spectrometry.

b) Recovery of DNA methylation in DNMT3B KO cells. Boxplots of relative methylation at H3K36me3 marked CGIs and all other CGIs methylated in HCT116 cells. P-values are from Wilcoxon rank sum tests. Lines=median; Box=25th–75th percentile; whiskers=1.5× interquartile range from box.

c) Recovery of DNA methylation in DNMT1 KO cells. Boxplots of relative methylation at H3K36me3 marked CGIs and all other CGIs methylated in HCT116 cells. P-values are from Wilcoxon rank sum tests. Lines=median; Box=25th–75th percentile; whiskers=1.5× interquartile range from box. The sample at day 28 failed quality control during processing.

**Supplementary Figure 3**

**Testing whether re-methylation following 5-aza-dC treatment can be explained by the outgrowth of cells escaping hypomethylation.** Schematic of the expected dynamics of re-methylation under two different scenarios, *de novo* methylation or the outgrowth of cells escaping hypomethylation due to a fitness advantage. If re-methylation was entirely explained by outgrowing cells, then the rate of re-methylation relative to the initial methylation level is expected to be identical for each CGI. The upper panel depicts a schematic of methylation levels at a single CGI in a population of cells over time under both models. The lower panel depicts the expected dynamics of re-methylation at different CGIs over time.

### Figure S1

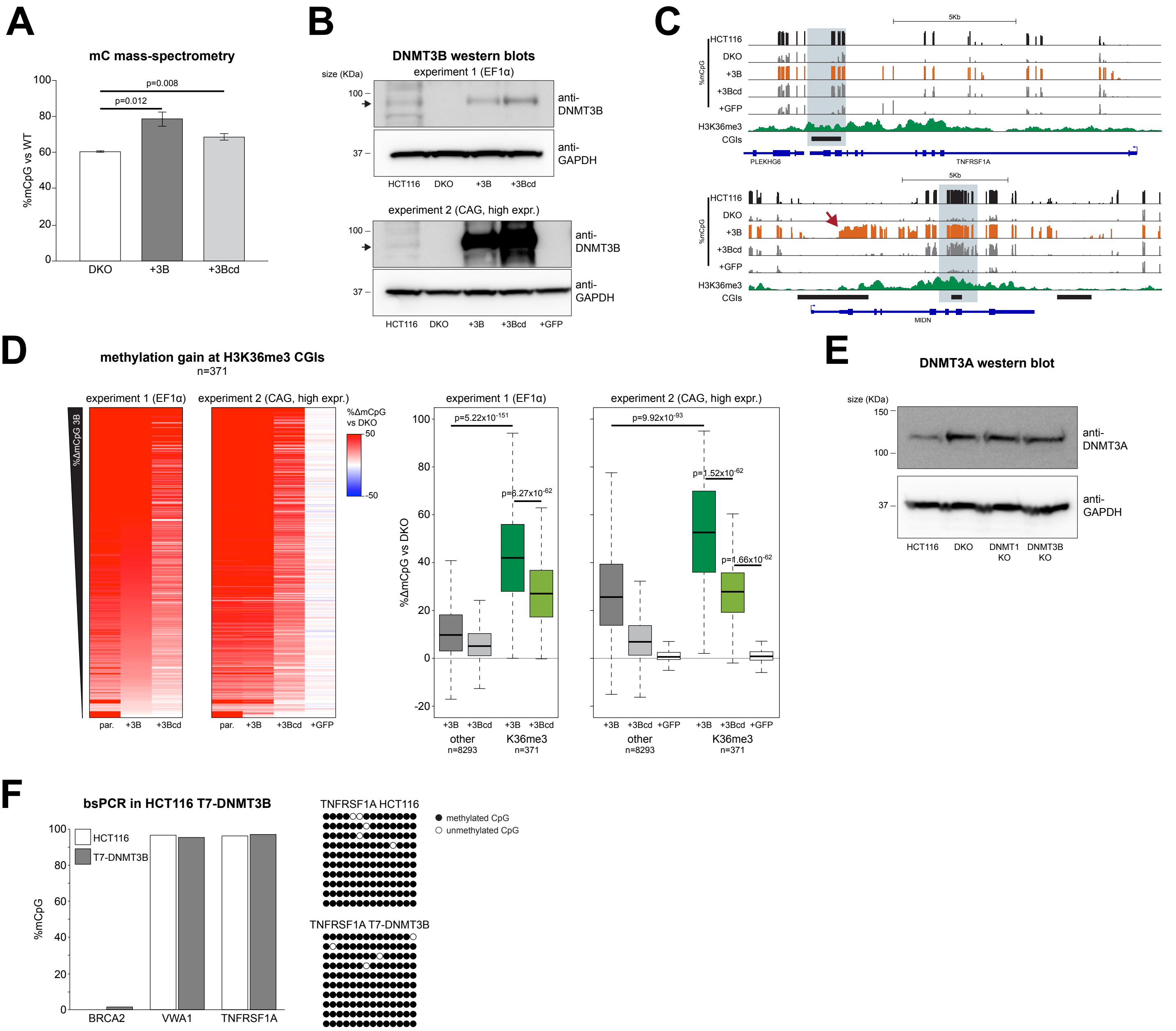

A
mC mass-spectrometry
p=0.008
p=0.012
%mCpG vs WT
100
80
60
40
20
0
DKO
+3B
+3Bcd
B
DNMT3B western blots
size (KDa)
experiment 1 (EF1α)
100
37
anti-DNMT3B
anti-GAPDH
HCT116
DKO
+3B
+3Bcd
experiment 2 (CAG, high expr.)
100
37
anti-DNMT3B
anti-GAPDH
HCT116
DKO
+3B
+3Bcd
+GFP
C
5Kb
HCT116
DKO
+3B
+3Bcd
+GFP
%mCpG
H3K36me3
CGIs
PLEKHG6
TNFRSF1A
5Kb
HCT116
DKO
+3B
+3Bcd
+GFP
%mCpG
H3K36me3
CGIs
MIDN
D
methylation gain at H3K36me3 CGIs
n=371
experiment 1 (EF1α)
experiment 2 (CAG, high expr.)
%ΔmCpG 3B
%ΔmCpG vs DKO
50
-50
par.
+3B
+3Bcd
par.
+3B
+3Bcd
+GFP
experiment 1 (EF1α)
experiment 2 (CAG, high expr.)
p=5.22x10$^{-151}$
p=6.27x10$^{-62}$
p=9.92x10$^{-93}$
p=1.52x10$^{-62}$
p=1.66x10$^{-62}$
%ΔmCpG vs DKO
100
80
60
40
20
0
-20
+3B
+3Bcd
+3B
+3Bcd
other
n=8293
K36me3
n=371
+3B
+3Bcd
+GFP
+3B
+3Bcd
+GFP
other
n=8293
K36me3
n=371
E
DNMT3A western blot
size (KDa)
150
100
37
anti-DNMT3A
anti-GAPDH
HCT116
DKO
DNMT1 KO
DNMT3B KO
F
bsPCR in HCT116 T7-DNMT3B
HCT116
T7-DNMT3B
%mCpG
100
80
60
40
20
0
BRCA2
VWA1
TNFRSF1A
TNFRSF1A HCT116
TNFRSF1A T7-DNMT3B
methylated CpG
unmethylated CpG

### Supp Figure 2

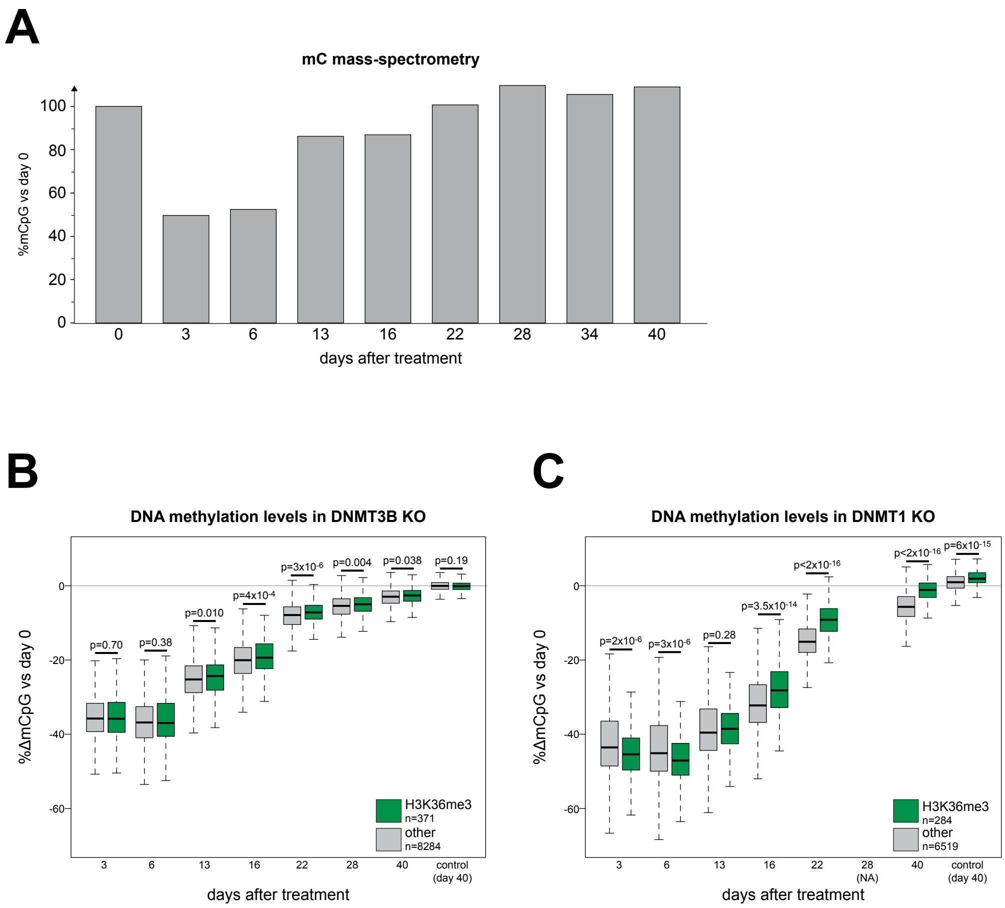

### Figure S3

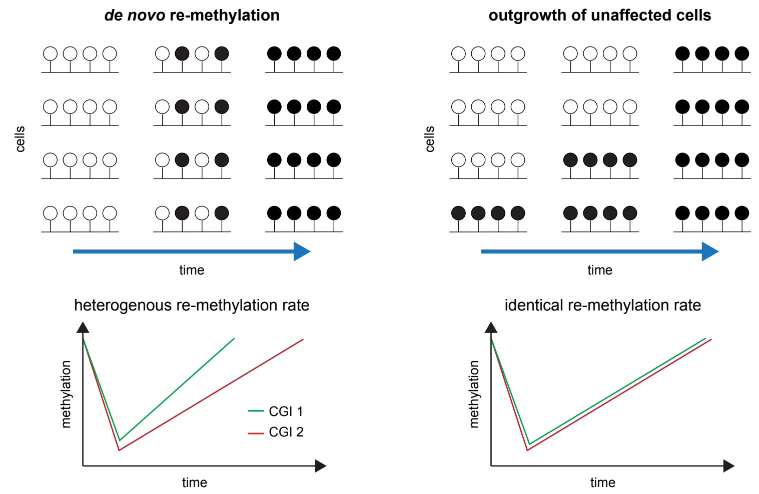
*de novo* re-methylation
outgrowth of unaffected cells
cells
time
heterogenous re-methylation rate
identical re-methylation rate
methylation
CGI 1
CGI 2
time
